# Tracing the regulatory atlas of non-coding RNA in human labour

**DOI:** 10.64898/2026.07.06.736857

**Authors:** Murali Aadhitya Magateshavren Saras, Shandar Ahmad, Roger Smith, Mithun K. Mitra, Sonika Tyagi

## Abstract

The early onset of labour increases mortality and developmental risks for a human newborn. Key genes in human labour have been investigated using multiple modalities, but their regulation by non-coding RNA (e.g. lncRNA and miRNA) remains incomplete. This study explores the three-way relationship between labour-associated transcription factors (TFs), miRNA and lncRNA suggested by the competing endogenous RNA (ceRNA) hypothesis, to understand the underlying regulatory framework. Experimentally validated miRNA-lncRNA interactions are modelled using five distinct machine learning (ML) architectures to predict 20469 labour-linked miRNA-lncRNA interactions. Known mRNA-ncRNA interactions from databases were included to construct a tripartite network, and a subset of 9989 labour-linked network motifs containing TFs were isolated and analysed. Gene enrichment of nodes in TF-lncRNA-miRNA network, as well as validation from public myometrial datasets indicate high significance in contractile pathways including immune signalling. Experimentally unconfirmed tripartite network motifs have been found, and we elaborate on their potential regulation in labour using 8 TF-lncRNA-miRNA network motifs. A unified ncRNA-TF regulatory atlas in labour has been synthesized, and a complete summary of the tripartite network motifs can be accessed and visualised using the user-friendly, public database.

## Introduction

Human myometrial smooth muscle cells (HMSMCs) partake in uterus contractions, called labour, to lead to childbirth. The onset of labour is crucial for a child being born at term (clinically defined as 37 weeks of gestation). Early onset and premature births increase a newborn’s risk of mortality and developing chronic disorders such as obesity, early onset heart disease and kidney failure [1]. Currently, the molecular mechanisms that regulate the onset of uterine contractions are not fully known, impeding the development of therapeutics to target preterm labour.

The contractions of HMSMCs are brought about by specific gene regulation through a hierarchical process stemming from hormone signalling [2]. During the late stages of pregnancy, progesterone levels increase while estriol and Corticotropin-releasing hormone (CRH) levels sharply rise [3, 4]. The signalling cascade affects transcription factors (TFs) from families such as the Activator Protein 1 (AP1), Nuclear factor kappa B (NF-*κ*B), Estrogen receptor (ER), and Progesterone receptor (PR) [5]. Additional TFs such as SMAD, HOXA, FOXO, MYB, ELF3 and SOXO have been validated to regulate the expression of labour-specific genes [6]. Thus, the gene expression trajectories in labouring samples change two-to-four weeks before delivery, indicating a transition from pregnancy maintenance to pre-labour biology [7]. Differential expression analysis of labouring and non-labouring myometrial samples have revealed key genes and non-coding RNA (ncRNA) responsible for pregnancy and contraction [8, 5, 2, 9, 10]. Our previous work identified three clusters of co-expressed genes linked to 15 TFs. The clusters were identified with specific functions: labouring samples had enriched immune signalling and matrix degradation activity, while non-labouring samples retained roles of muscle quiescence and embryonic development [2].

Gene expression and regulation by long ncRNA (lncRNA) and microRNA (miRNA) has been suggested across various studies, and the competing endogenous RNA (ceRNA) hypothesis suggests an additional lncRNA-miRNA interaction to form a tripartite regulation [11, 12, 13, 14, 15, 16]. lncRNAs co-expressed with proximal genes are found to sequester miRNA and control miRNA-mediated gene silencing in cancer and ischemic stroke [13, 14, 15, 16]. ncRNA-dependent gene regulation in labour and ncRNA-TF co-regulation has been studied, albeit for a few cases [10, 17, 18, 19]. Members of the miR-200 family facilitate labour by repressing ZEB1 and ZEB2 [20], and lncRNAs UCA1, XIST and MIR200CHG are linked to the expression levels of miR-203 and miR-141 [21, 22, 11]. In our previous work, we have identified multiple ncRNAs (15 lncRNAs, 27 miRNAs) collective with the co-expressed gene clusters and associated TFs, and we expect them to be a part of the ceRNA-based regulation in labour [2, 23]. Currently, evidence supporting lncRNA-miRNA links in labour pathways can only be inferred from other studies. A comprehensive atlas describing the global regulatory network between miRNA, lncRNA and genes (including TFs) in labour is lacking.

In this study, we use machine learning (ML) methods to predict potential interactions between differentially expressed lncRNA and miRNA in human labour samples. These ML methods were trained on experimentally validated interactions between miRNA and lncRNA to model direct sequence based interactions. mRNA interaction information from databases were integrated with the predicted ncRNA to construct a network and identify tripartite relationships. TFs within the network and their motifs are identified and annotated to gain insight into the regulatory relationships. Thus, our work fills a critical knowledge gap on the regulatory interactions between lncRNA-miRNA and associated TFs in human labour.

## Results

In this section, we describe the ML methods used, their performance, and a summary of the lncRNA-miRNA interactions predicted. We construct a tripartite network, and gene ontology enrichment of the network nodes indicate functional roles in labour-associated pathways. We summarise our investigations on experimentally validated as well as on unconfirmed interactions using literature evidence and public omics datasets.

### Labour-linked miRNA-lncRNA interactions predicted using five Hybrid ML models

Experimentally validated lncRNA-miRNA interactions with their sequences were modelled using five different, hybrid ML architectures, and the best-performing model metrics are depicted in Figure 1A. Among all the ML tools, the highest validation metrics were obtained using the TEC-LncMir model (Accuracy - 0.989, F1 - 0.988), followed by SPGNN (Accuracy - 0.971). PMLIPred and the ensemble method had similar accuracy scores of 0.871 and 0.863, respectively. The best ensemble model metric with an accuracy score of 0.86 comprised four models - the RF modelled on feature categories 1 and 2 and the XGB modelled using categories 2 and 4 (See Methods 6.4, Table 2). The best-trained model of genomicBERT achieved an accuracy score of 0.69. As an LLM model, the small size of training data is suspected to have affected the predictive power of this model.

**Fig. 1.**
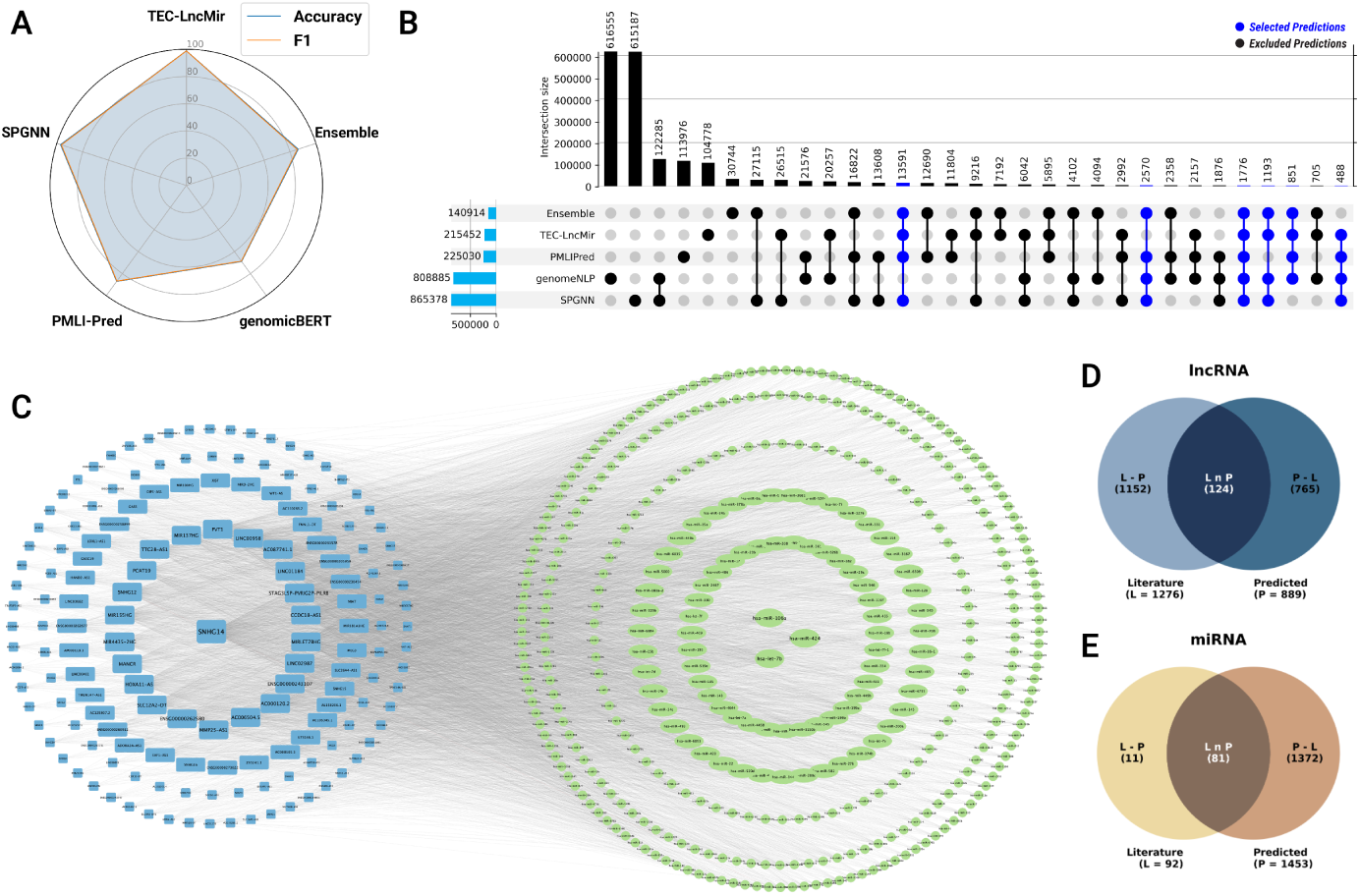
A) Radar plot showing the model accuracies and F1 scores achieved on the training data. The range is from 0-100; B) UpSet plot showing the modelling results and the overlap of lncRNA-miRNA predictions from 5 methods. The horizontal bars in the lower left indicates the total number of positive predictions given by each model. The vertical bars indicate the overlap of results across the different models, indicated by the dark, connected circles. C) Cytoscape network of lncRNA-miRNA interaction predictions from 4 and more concordant model results. D,E) Venn diagram depicting lncRNA and miRNA common between literature sources and predicted results.

Using the trained models, the test dataset interactions were predicted and the SoftMax scores were rounded off by a threshold to collect the positive predictions. The UpSet plot (Figure 1B) depicts the overlapping results across the 5 models. The SPGNN model had the highest number of positive predictions (n=865738), followed by genomicBERT (n=808885). However, genomicBERT had the highest number of unique predictions unseen in other models (n=616555). The other models see a stark decrease in the total positive predictions from the test dataset, but retained higher numbers of unique predictions. Each ML model has advantages owing to the architecture and feature representation methods, and hence, a consensus result from four or more models was used for network analysis.

Based on this criterion, 20469 lncRNA-miRNA interactions were selected for downstream analysis (Figure 1B: highlighted in blue). The bipartite interaction predictions (n=20469) compiled from the models consisted of 366 unique lncRNA and 1124 unique miRNA (Figure 1C). However, only 14% of lncRNAs (n=124) and 5% of miRNAs (n=81) are reported in literature pertaining to labour (Figure 1D, 1E). The predicted list of lncRNA-miRNA interactions (n=20469) was combined with miRNA-mRNA interaction (n=800756), and lncRNA-mRNA (n=152979) pairs. A total of 106,455 tripartite network motifs (3-node cliques) comprising 95 lncRNA, 428 miRNA and 3937 mRNA (inclusive of 369 TFs) were identified using the networkX package in Python. TF-specific tripartite links were isolated (n=9989), and gene ontology enrichment results of each type is discussed in the next section.

### GO enrichment of predicted ncRNA and associated TFs suggest contractile regulation in HMSMCs

Gene enrichment was carried out independently for each class of RNA: TFs with STRING, lncRNAs with g:Profiler, and miRNAs with MiEAA.

The enrichment from 367 miRNAs found within TF-linked network motifs indicates localisation in placental and umbilical tissues (Figure 2A,B). Here, 33 miRNAs have been identified to be placenta-specific, and 104 miRNAs are indicated as cerebrospinal fluid-specific (CSF) localisation. Placenta-derived miRNAs such as MIR155, MIR653, and MIR618 have been correlated with pathological effects during pregnancy [24]. CSF-localised MIR1244 and MIR452 are upregulated and downregulated, respectively, and their roles in progesterone regulation has been validated [25, 26]. Figure 2A illustrates that gene regulatory functions affect pathways related to HMSMC transition and immune response. The relationship of the motif miRNA to labour is emphasised by pathways such as myometrial relaxation and contractile pathways, eicosanoid synthesis, and prolactin signalling. Additional evidence is supported by the terms ‘TGF*β* signalling’, ‘NF-*κ*B signalling’, ‘IL-1 and IL-4 signalling’, indicating towards the activation and regulation of immune response.

**Fig. 2.**
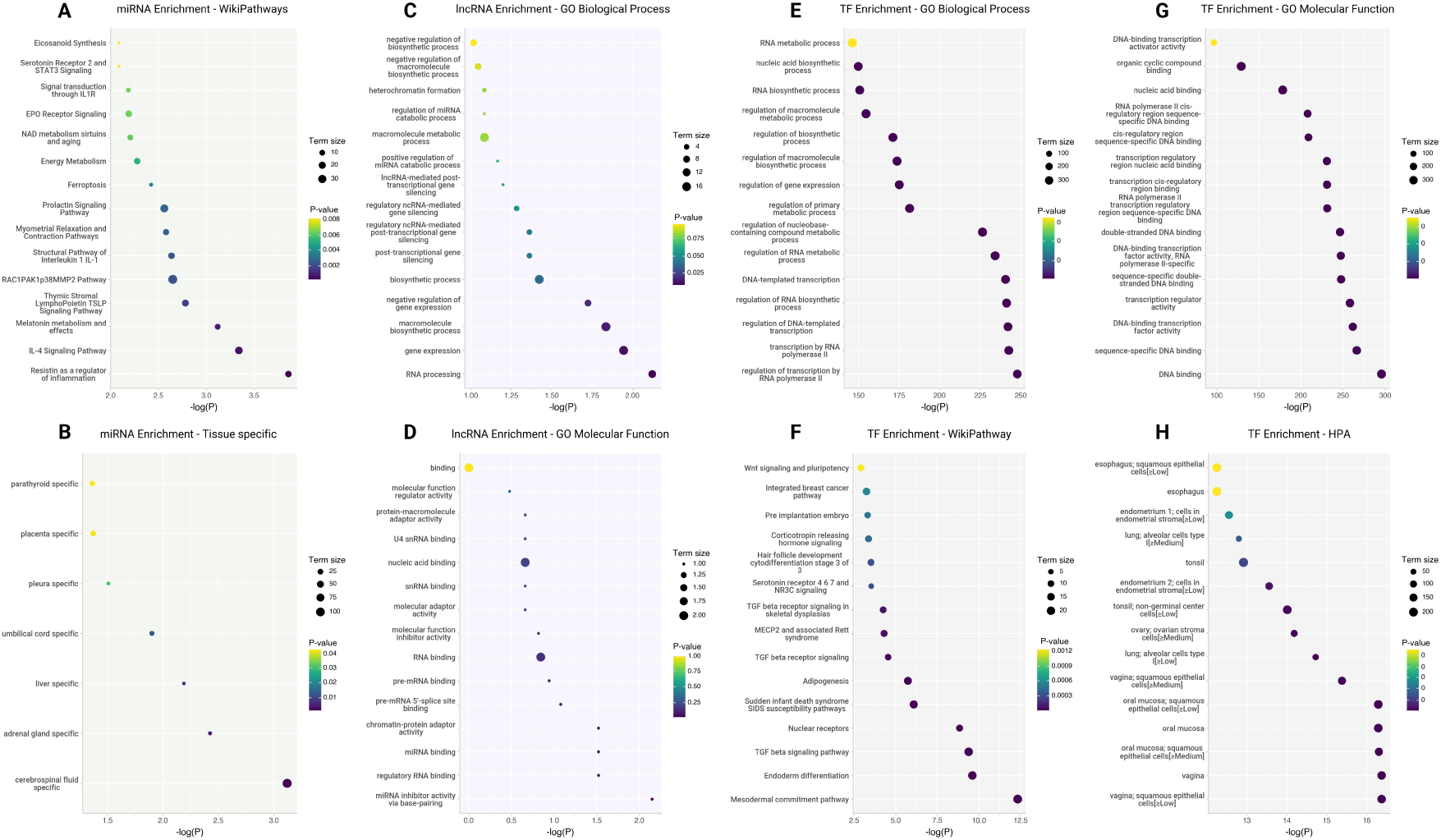
Gene Ontology enrichment of miRNA (A,B), lncRNA (C,D) and TFs (E-H) compiled from tripartite network motifs.

Gene enrichment and ontology annotation for 42 lncRNA nodes using g:Profiler [27] indicate high similarity with miRNA enrichment, post-transcriptional gene silencing and regulation of miRNA catabolic process (Figure 2C,D). The functional annotation indicates that the network motif lncRNA predominantly possesses RNA-binding activity with an inhibitory function (Figure 2C). Few lncRNAs such as NEAT1 and PVT1 have been annotated with ‘miRNA binding’, ‘RISC complex binding’ and ‘regulatory ncRNA-mediated gene silencing’, indicating ceRNA-style tripartite regulation.

The gene ontology:biological processes (GO:BP) for the list of TFs has yielded terms linking them to their function in transcriptional regulation through binding to RNA polymerases, DNA and RNA (Figure 2E,G). Hormone signalling pathways and their cascade systems are enriched, such as the estrogen receptor pathway and the CRH signalling pathway (Figure 2F). The localisation of the TFs show enriched results in vagina, endometrium and ovarian stromal cells (Figure 2H). While CRH is commonly known for inflammatory responses, placental CRH synthesis is considered to regulate fetal maturation and the onset of labour [28, 29]. TGF*β* plays a central role in labour onset by contributing to the remodelling of the myometrium, placental development and immune tolerance. TGF*β* binds and activates the SMAD receptors, which form complexes to initiate transcription and contribute to HMSMC contractions [30]. TGF*β*, as a cytokine, also affects the immune tolerance during pregnancy [31]. Further, increased adipogenesis prior to parturition increases the fat cells of the fetus in preparation for the extrauterine conditions after birth.

The enrichment results of the predicted miRNA-lncRNA links limited to TF associations reveal labour-associated regulation and molecular function. We investigate and report on the linked pathways between ncRNA and TFs in the next section.

### Network Motifs reveal published and potential interactions linked to labour

All the motif networks were cross-referenced to the information collected from databases. 6969 network motifs were identified without any experimental evidence (Supplementary Table S2). We prioritised exploring the motif sub-networks of 35 TFs identified as potential regulators from a previous myometrial analysis [2], and 520 tripartite links were filtered which involved 14 of these 35 TFs. Out of 520, we were able to find experimental evidence for 175 tripartite links. We illustrate the nature of the experimental evidence for a few representative motifs below:

### NEAT1/MIR485/STAT6

There is indirect evidence of the tripartite relationship between STAT6, NEAT1 and miR485. Huang *et al*. mention NEAT1 increasing expression of STAT6 to promote cytokine expression [32], and Xu *et al.* describe the ceRNA relationship between NEAT1 and MIR485 [33]. The sponging effect of NEAT1/miR485 could be the mechanism for the potential downstream effect on STAT6 expression.

### GAS5/MIR145/FOXO3, GAS5/MIR485/FOXO3, GAS5/MIR23a/FOXO3

Direct GAS5-mediated regulation of FOXO3 by sponging MIR182-5p has been validated [34]. Additional ceRNA based evidence between GAS5 and FOXO3 can be linked with MIR145 [35, 36, 37], MIR485 [38, 39], and MIR23a [40, 41, 42] across different phenotypes and tissues.

### MIR155HG/MIR454/MYB

MYB physically associates with the promoter of MIR155HG and stimulates its transcription [43], and MIR454 has been experimentally validated to be linked to MYB [44]. MIR454 and MIR155 (part of MIR155HG) have ben shown to be significantly elevated in peripheral Blood of Uveal Malignant Melanoma Patients [45].

### SMAD5-AS1/MIR106-5p/SMAD5

Zheng *et al.* outline the sponging effect of SMAD5-AS1 on MIR106-5p to affect SMAD5 activity [46]. An indirect regulation in labour can be inferred based on the Epithelial to Mesenchymal transition pathways (EMT) targeted by SMAD5. The investigation was demonstrated in nasopharyngeal carcinoma, but their focus on Epithelial to Mesenchymal transition pathways (EMT) is also present in myometrial samples during labour. This suggests that the regulation of SMAD5 could be playing a role during labour as well. The remaining 345 network motifs lacked experimental evidence from databases. However, we hypothesise that they might be potential novel ceRNA links worth investigating for their roles in labour: CHANGE FLOW- ADD that we are Speculating the regulation between the 375 lists, and we have explored on a few representative plots

### LINC-PINT/MIR27B/RUNX1 and LINC-PINT/MIR27B/NFE2L2

MIR27B is found to directly regulate RUNX1 and NFE2L2 in cancer studies to suppress cell migration and EMT [47, 48, 49]. LINC-PINT controls cell proliferation and EMT through ceRNA regulation [50]. We propose that the LINC-PINT/MIR27b regulation is an undiscovered link that contributes to HMSMC transition.

### SLC7A11-AS1/MIR5590/NFE2L2 and SLC7A11-AS1/MIR106A/NFE2L2

NFE2L2 is a key transcription factor in the cellular defence against oxidative stress, playing a crucial role in cancer cell survival [51]. In colorectal cancer, SLC7A11-AS1 overexpressed tissues upregulated SLC7A11 and NFE2L2 [52]. MIR5590 and MIR106A both restrict tumour growth by repressing cell proliferation [53, 54]. Our results indicate that MIR5590 and MIR106A are potential undiscovered links, binding the co-regulation between NFE2L2 and SLC7A11-AS1.

### GAS5/MIR503/FOXO3, GAS5/MIR516B/FOXO3, GAS5/MIR331/FOXO3

ceRNA regulation of FOXO3 by GAS5 has been linked to multiple miRNAs: MIR182 and MIR9 [34, 12]. Our results highlight additional miRNAs such as MIR503, MIR331 and MIR516B to be involved in the ceRNA regulation.

### DLGAP1-AS2/MIR2114/ESR1

ESR1 mRNA expression levels are directly correlated with DLGAP1-AS2. ESR1 was significantly increased in DLGAP1-AS2-overexpressing cells but dramatically reduced in DLGAP1-AS2 knockdown cells [55]. Current literature links the correlated expression based on another mRNA AFF3, but our results indicate that MIR2114 could be another regulatory mechanism connecting DLGAP1-AS2 and ESR1 expression.

We have explored only on a subset of the unconfirmed interactions, and an investigation is needed to validate all potential interactions.

Validation of network nodes from public myometrial labouring datasets

#### TF found as DEGs in Labour are linked with 421 tripartite motifs

Public datasets from myometrial term and preterm sources with contrasting labouring and non-labouring phenotype were identified and analysed for differentially expressed genes (DEGs) (Figure 8). Using the meta-analysis approach, 118 DEGs from term samples, and 406 from preterm studies were identified, with 72 DEGs found common (Figure 3A, Supplementary Table S3).

**Fig. 3.**
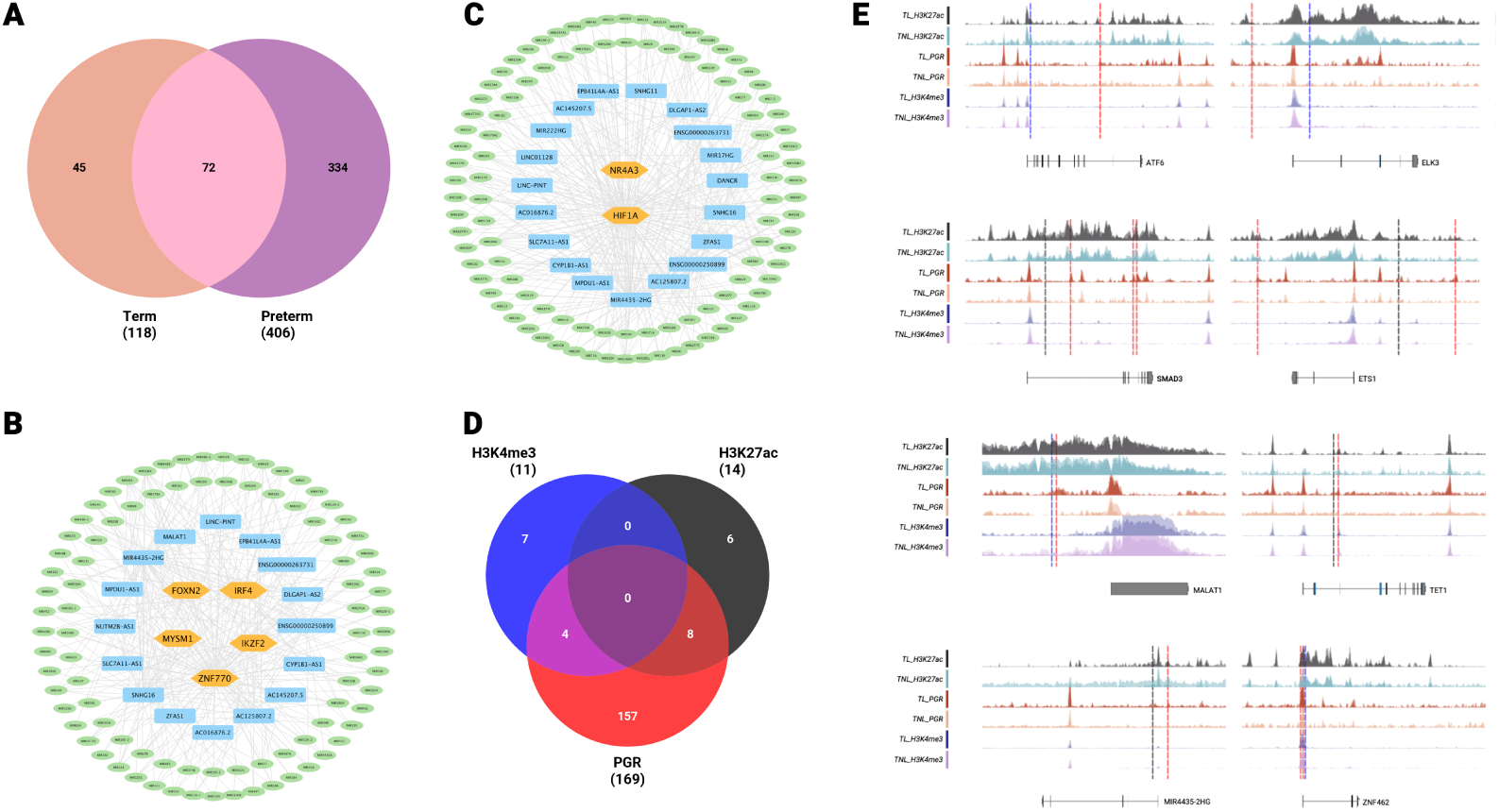
A) RNAseq DEGs identified from term and preterm myometrial samples; Motif networks identified with meta-analysis DEGs - Preterm (B), Term (C); D) Differential binding results of ChIPseq data - PGR, H3K4me3, H3K27ac; E) Track figures of genes with common differential binding sites; Vertical dotted lines correspond to differential binding sites of PGR (Red), H3K4me3 (Blue), H3K27ac (Black).

We found that two TFs identified as DEG within term samples were also a part of the tripartite network. NR4A3 and HIF1A form 223 tripartite motifs with 19 unique lncRNA and 104 unique miRNA (Figure 3C). From preterm labouring samples, 406 DEGs were identified but only 5 TFs (IRF4, FOXN2, MYSM1, ZNF770, IKZF2) were common with the tripartite network. A total of 198 motifs linked to 16 lncRNAs and 90 miRNAs were mapped to those 5 TFs and visualised (Figure 3D). Although there were no common TFs found within the motif-DEGs between term and preterm, 14 lncRNAs and 38 miRNAs were present in common within the shortlisted networks.

#### Differential PGR binding observed in 169 motif nodes suggest HMSMC transition

ChIP-seq differential binding analysis was performed using the public dataset GSE202027, containing term samples with contrasting labouring and non-labouring phenotype. The annotated differential binding analysis from all available proteins and histone marks (PGR, H3K4me3 and H3K27ac) were overlapped with the tripartite motif nodes, and visualised in Figure 3B, 3E (Supplementary Table S4). H3K4me3 as an active promoter mark may suggest transcription initiation regions, while H3K27ac highlights enhancer activity.

Differential PGR binding overlapped with 169 network nodes (109 TFs, 7 lncRNA and 53 miRNA), while H3K4me3 binding had 11 nodes (8 TFs, 1 lncRNA and 2 miRNA), and H3K27ac was found to overlap with 14 nodes (7 TFs, 1 lncRNA and 6 miRNA). PGR-specific binding enriched roles in adipogenesis, mesodermal commitment pathway and TGF*β* signalling, suggesting HMSMC transition. However, differentially bound TFs from H3K27ac and H3K4me3 do not show any enrichment.

There were no common nodes found with all 3 differential peaks, but a few are found with overlapping peaks between PGR-H3K4me3 (3 TFs, 1 lncRNA) and PGR-H3K27ac (4 TFs, 1 lncRNA, 3 miRNA) (Figure 3B, 3E). The common nodes between PGR-H3K4me3 are 3 TFS: ATF6, ELK3, ZNF462, and one lncRNA: MALAT1. ATF6 plays a critical role in the negative regulation of placental growth factor (PlGF), to manage protein folding and in maintaining endoplasmic reticulum stress [56, 57]. MALAT1 and ELK3 in association with other genes are involved in trophoblast invasion and migration [58]. ZNF462 is not cited with a direct role in labour, but a loss-of-function affects prenatal development and the course of pregnancy and birth[59].

Between H3K27ac-PGR binding sites, 4 TFs (ETS1, SMAD3, TET1, TFAP2C), 1 lncRNA (MIR4435-2HG) and 3 miRNAs (MIR1294, MIR326, MIR493) were found in common with the network motifs. The roles of the identified nodes are highly cross-linked with labour: ETS1 is involved in extra-cellular matrix (ECM) remodelling, while TET1, MIR326 and TFAP2C regulate trophoblast growth, migration and invasion[60, 61, 46, 62]. lncRNA MIR4435-2HG plays a role in epithelial-to-mesenchymal transition (EMT), reducing cell adhesion and shifting to an apoptic state during labour[63, 64]. SMAD3 is associated with increased production of proinflammatory and prolabour mediators in the human myometrium[46, 30]. MIR493 in association with SNHG14 is related to adverse pregnancy outcomes in people with diabetes mellitus[65]. MIR1294 has not been found with any evidence to labour or associated risks yet.

The evidence suggested by the motif overlap along with PGR binding and the histone marks indicates the regulation and activation of key genes with critical roles in labour.

#### Network nodes are localised in specific chromosomes

The localisation of network motif nodes within the human genome reveal that miRNAs, lncRNAs and TFs are unevenly located across chromosomes. The ratio between observed and expected frequencies for each RNA type is visualised in Figure 4A.

**Fig. 4.**
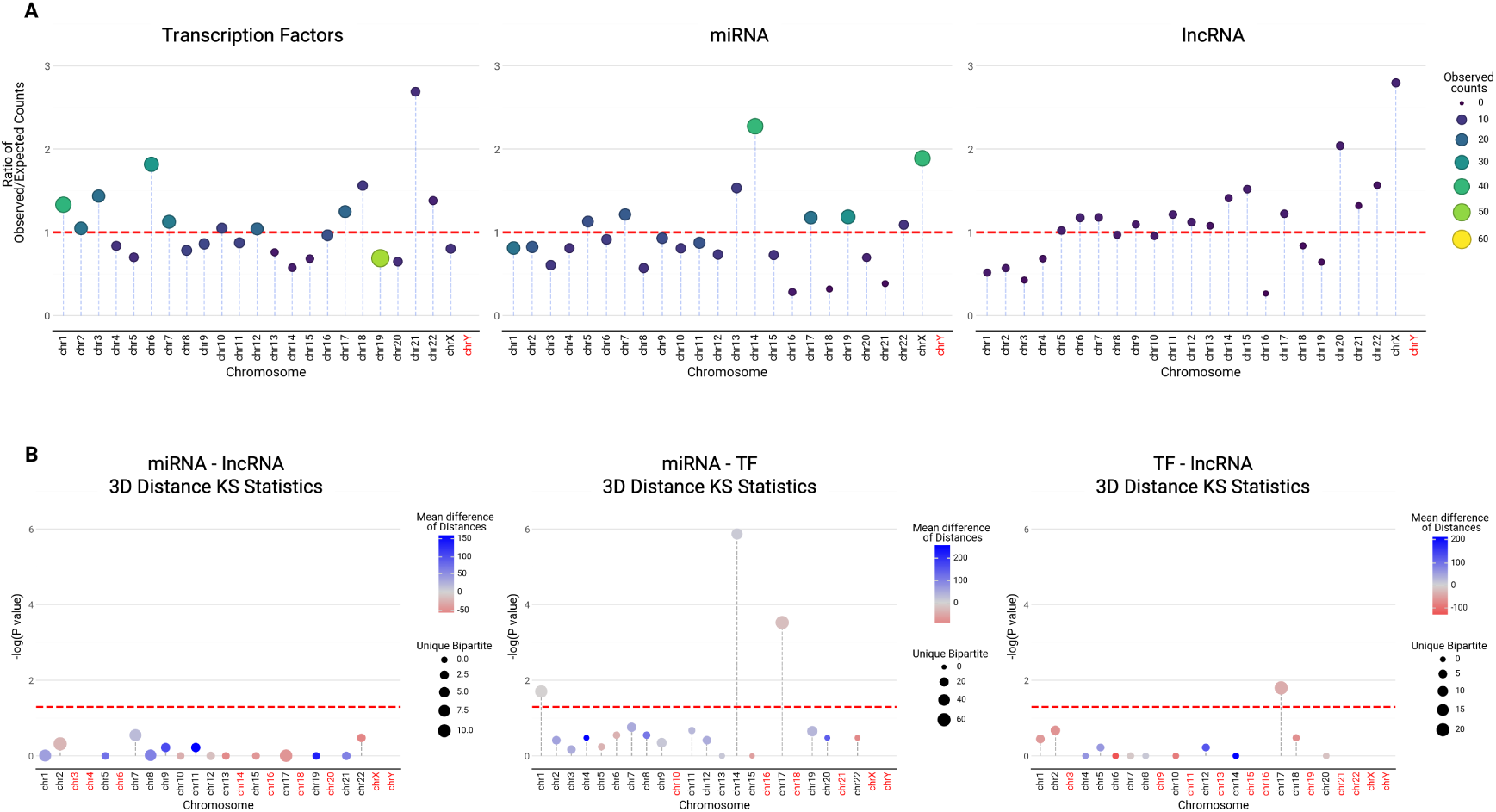
A) Chromosome-specific observed vs expected frequency ratios of miRNA, TFs and lncRNA found in the tripartite network. The observed frequencies are represented by both bubble size and colour, and the red-dotted line marks ratio = 1; B) Chromosome vs -log(p-value) calculated using Kolmogorov-Smirnov 2-sided test on observed vs expected distances. The number of bipartite combinations used for statistical testing is represented as the bubble size, and mean difference in distance is shown in the colour gradient. The red-dotted line signifies p-value=0.05. The chromosome names in red indicate lack of data for statistical testing.

The observed frequencies compared against their expected frequencies help us understand the relative abundance of each RNA type involved in labour regulation from each chromosome. We notice that the patterns are irregular across the 3 types: miRNA, lncRNA and TFs. miRNA from the motif network are prevalent in chromosomes 14, X, 19 and 17 in decreasing order, and their ratios reflect a similar pattern. However, TFs and lncRNA depict patterns unlike the ratio seen in miRNA. Chromosome 19 has the highest number of mapped TFs (n=53), but the ratio is less than 1, suggesting that only a small subset of the TFs in Chromosome 19 are involved in labour regulation. Chromosomes 21 and 22 show a higher ratio despite low counts hinting that their role in labour is enhanced. ETS2, ATF4, GABPA, and MAFF are TFs found within chromosomes 21 and 22, and their ontology analysis show their roles in ‘Myometrial relaxation and contraction pathways’ and ‘NFE2L2 regulating anti-oxidant/detoxification enzymes’. The NFE2L2 regulation helps to combat oxidative stress, which plays a crucial role in labour [66, 67].

Similarly, lncRNAs such as DSCAM-AS1, XIST, MIR222HG, SNHG11, SNHG17 localised in chromosomes 20, 21, 22 and X have the highest ratio, hinting at their activity in labour regulation.

In the following section, we investigate if the network motif nodes are also co-localised within the same TAD boundaries and summarize on their regulatory links.

#### Network motifs indicate inter- and intra-TAD relationships

The motif nodes were mapped to the TAD regions and 3D distances calculated using the myometrial samples from GSE244735. A total of 1753 TAD regions, ranging from 400 Kb - 3 MB were identified using a 100 KB resolution and a sliding window size of 500 KB across all chromosomes (Supplementary Figure S5). The network motif nodes, mapped to their genomic loci, was overlapped with the TAD boundaries to identify co-occurring motifs. However, no TAD regions were identified where all 3 types of motif nodes (TF, miRNA, lncRNA) co-occurred. A bipartite mapping analysis was carried out (TF-lncRNA, lncRNA-miRNA, miRNA-TF) and 16 combinations (2 lncRNA-TF, 14 TF-miRNA) were found localised within the same TAD region across 8 chromosomes (Supplementary Table S5).

Additionally, 3D distances were calculated between motif nodes found within the same chromosome to investigate cis-regulatory relationships. The statistical analysis between motif distances and expected distances reveal significance in specific chromosomes for each category. Between TF-miRNA pairs, 3 chromosomes with significant differences in distances were identified (Figure 4). TF-miRNA average observed distances were found to be lower than expected in Chromosomes 1 and 17, while they are higher than expected in Chromosome 14. The lncRNA-TF category identified only chromosome 17 with a significant difference between observed and expected distances. Between miRNA-lncRNA combinations, the distances are varied across chromosomes, but none have significant separation.

## Discussion

### Regulatory Tripartite Motifs identified for labour-associated pathways

Recent advances in ML have brought new methods to predict miRNA-lncRNA interactions [68, 69]. In this study, we employed five different ML tools to predict and compile miRNA-lncRNA interactions specific to labour. We identified 20469 predictions supported by four and more models used in this study.

Compiling the miRNAs, we found that a major list of interactions overlap with SNHG14 (1108), PVT1 (426) and CCDC18-AS1 (345) lncRNAs. SNHG14 has been identified as a cancer-associated lncRNA, and the sponging effect with multiple miRNAs have been reported across the literature. The role of PVT1 in preeclampsia is studied, but not yet explored in the context of labour, and CCDC18-AS1 in unknown in labour[2, 70]. We find that a majority of the miRNAs are involved directly and indirectly in labour through multiple pathways such as cell transport and silencing via complexes. A comparison to the evidence collected from the databases (see Methods) indicates 6048 (∼30%) bipartite miRNA-lncRNA links, out of 20469 predicted, to be experimentally validated by prior studies. The complete predicted list was augmented with mRNA-ncRNA bipartite links, and 9989 TF-specific interactions were analysed further. Gene enrichment of lncRNA, miRNA and TFs indicate roles in labour-linked pathways: Progesterone regulation and TGF*β* signalling by miRNAs, RNA-binding and inhibitory functions of lncRNAs with evidence suggesting their regulation of labour-associated genes, and TFs involved in estrogen receptor pathways and CRH signalling pathways. The network motifs nodes are also found localised in endometrium, placenta and ovarian stromal cells.

Out of a total 9989 TF-lncRNA-miRNA network motifs, 6969 tripartite links could not be matched to the experimentally validated list from multiple databases. An exploratory investigation to cross-reference matched and unmatched network motifs was performed after shortlisting based on 35 TFs identified from our previous myometrial analysis [2]. Network motifs overlapped with 14 TFs, and out of 520 shortlisted tripartite interactions, 175 of them matched with experimentally validated data. A literature survey and analysis revealed direct and indirect evidences for the network motifs. SMAD5-AS1 levels control EMT, cell proliferation, migration, and invasion by sponging MIR106a and regulating SMAD5. Similarly, GAS5-FOXO3 ceRNA-based regulation has been reported in multiple cancer types [34, 12, 71, 41]. Additionally, indirect but linked evidence has been found for a few of the network motifs. For example, LINC-PINT inhibits EMT and cell proliferation [50], while MIR27B also regulates RUNX1 to control cell migration [47]. We hypothesize that this is a missing regulatory link affecting EMT, despite the lack of direct experimental/literature-based evidence between LINC-PINT and MIR27B. Similarly, ESR1 and DLGAP1-AS2 are co-expressed, and our results indicate that MIR2114 could be the third component, forming a ceRNA regulation.

We present our results of an interconnected network of tripartite links with TFs-lncRNA-miRNA. While only a subset of the tripartite network motifs are found with experimental evidence, the gene enrichment results suggest more ncRNAs contributing to labour. Thus, the network serves as a foundation to expand research to understand regulation in labour.

### Myometrial Omics data inference

We utilised public, human, myometrial omics datasets to validate the tripartite network motifs identified in this study.

Differentially expressed genes from term and preterm myometrial datasets between labouring vs non-labouring phenotypes had a small overlap with the nodes of the network, and none of the lncRNAs or miRNAs were common. The low overlapping numbers (2 TFs from term and 5 TFs from preterm) might be due to the lack of a comprehensive list of experimentally validated interaction data, or the threshold of 4+ model results used for network construction. ChIP-seq analysis was performed on term labouring vs non-labouring phenotype, and PGR differential binding analysis overlapped with 169 nodes from the tripartite network (TFs = 109, miRNAs = 53, lncRNAs = 7). In comparison with RNAseq DEG data, only 18 genes (including 3 TFs) were found in common that are also differentially bound by PGR. Multiple prior literature report that accessible chromatin regions are not always linked to DEGs[72, 73], even within matched samples. Despite the experimental differences, we also observed peaks by histone marks H3K4me3 and H3K27ac, overlapping with the RNAseq data and the network motifs. Albeit partial, this validates that the network nodes are targeted by enhancers, promoters and TFs in labouring samples, and supports that the ncRNA-TF regulatory circuit is transcriptionally active. Furthermore, this strengthens evidence that the network motifs are not merely correlation-driven, but mechanistically connected. Thus, overlapping ceRNA tripartite motifs with histone ChIP-seq and protein ChIP-seq data serves to experimentally contextualise computationally inferred regulatory networks, identify active chromatin-associated regulatory circuits, prioritise biologically meaningful motifs, and uncover integrated epigenetic–transcriptional–post-transcriptional mechanisms.

We also investigated the myometrial landscape using a public HiC dataset to understand TAD localisation of the motifs. TAD regions limit promoter-enhancer interactions, and genes within a TAD can be co-expressed. We investigated the co-localisation of the network motifs, and no TAD regions were identified that contained all 3 nodes (TF, lncRNA, miRNA) of a predicted motif, even with sizes up to 3 MB. This aligns along the ceRNA hypothesis that if a competing element is present in proximity to the linked genes, then the competing nature will be rendered ineffective if all the gene-lncRNA-miRNA combinations are co-expressed. However, 16 bipartite links (2 lncRNA-TF, 14 TF-miRNA) were found co-localised within TAD regions across 8 chromosomes (Supplementary Table S5). We probed the 3D distances between bipartite combinations within the network, and found different patterns across chromosomes. Only a few chromosomes had significant difference in distances between miRNA-TF and lncRNA-TF combinations. No significant difference in 3D distance between motif nodes and expected values was observed for miRNA-lncRNA motifs across any chromosome. The relative scarcity of intra-TAD co-localisation and limited evidence of short-range cis interactions suggest that many motif components may function through inter-TAD or long-range trans-regulatory mechanisms. These findings indicate that labour-associated regulatory circuits in the myometrium may involve genomically distributed chromatin interactions. Further investigation with matched experiment will be necessary.

### Indication of a multi-regulatory Framework

It is interesting to note that 35,909 lncRNA and 2656 miRNA sequences were used for predicting, and the consensus across model predictions filtered 1124 miRNAs but only 366 lncRNAs. Constructing the tripartite network and isolating TFs shortlisted 42 lncRNAs and 370 miRNAs, indicating that only a few non-coding RNA partake in regulation.

The TFs HIF1A, FOXN2, IRF4, and NR4A3 are found common in RNAseq DEGs and within the tripartite network. Literature reports their significance in the onset of labour and potential risks to preterm birth through the regulation of labour- and immune-associated genes such as GJA1, CX43, PTGS2, PTGFR, and IL6[74, 75, 76, 77, 78, 79]. Although not identified as DEGs, PTGS2, PTGFR, and IL6ST are found within the tripartite network, and the associated network-ncRNA form a subset of the ncRNA linked to HIF1A and NR4A3. This suggests that co-regulation and co-expression of genes reported in literature is due to the regulation brought about by ncRNA elements. Incidentally, while no TFs overlapped between term and preterm meta-analysis, the ncRNA network was common with 14 lncRNAs (out of 16 and 19 unique lncRNAs). This reinforces that the regulatory framework of various genes might be controlled by a small subset of lncRNAs.

The complexity of ceRNA networks increase when a few non-coding elements are found common to a large number of genes, and this expands the concept of ceRNA to include multi-level co-regulation of genes by a small group of ncRNA. While our finding are brief and limited by the datasets, a deeper investigation is needed of the tripartite network to understand gene regulation in labour.

### Limitations and Future Study

The analysis and the results of this study are limited to the AGO-CLIP lncRNA-miRNA interactions curated from public databases. This was done to limit the dataset to a high-confidence experimental validation to ensure robustness in modelling. Additionally, the lncRNA-mRNA interactions were very few in number to complement the tripartite network creation. However, the expansion of the inclusion criteria could increase the sample size, and thus, the modelling efficiency. The databases used for gene enrichment and ontology mapping were biased towards cancer-based datasets, and hence the enrichment results of the tripartite network nodes were skewed towards cancer localisation and cancer-associated functions. We also note the limited, unmatched samples of ChIP-seq and HiC myometrial datasets used for validation.

## Conclusion

Various independent studies have investigated the molecular landscape of labour through different aspects: RNAseq, ChIPseq, ATACseq and HiC. Although each study has presented varying levels of information, the concept of regulation by ncRNA was not well elucidated. In this study, we employed multiple ML architectures to predict ncRNA relationships between known non-coding components in labour, and extended it to genes and transcription factors and identified regulatory relationships. The predicted networks and linked nodes have been validated using literature and myometrial public datasets on Microarray, RNAseq, ChIPseq and HiC. We present results of validated and potential regulatory links between TFs, lncRNA and miRNA in labour as a publicly accessible resource (https://gitlab.com/tyagilab/LMI_labour, Supplementary Figure 6).

## Declarations

### Code and Data Availability

All the codes used in this study can be found in the respective publications [80, 81, 82, 83]. The website link and the code is available at https://gitlab.com/tyagilab/LMI_labour.

### Author contributions statement

**MAMS-** Formal analysis, Visualization, Writing - Original Draft; **ST-** Conceptualization, Supervision, Writing - Review & Editing; **MKM-** Supervision, Writing - Review & Editing; **SA-** Writing - Review & Editing; **RS-** Writing - Review & Editing;

## Acknowledgments

We acknowledge Naima Vahab for her assistance in modelling using genomeNLP. We also acknowledge Yashpal Ramakrishnaiah, Jayden Wade, Evie Lockhart, Elizabeth Pollard, Maria Alshaikh, Bao Ho for working on the database.

## Appendix

## Abbreviations

HMSMC: Human Myometrial Smooth Muscle Cells
TF: Transcription Factors
RNA: Ribonucleic Acid
ncRNA: non-coding RNA
ceRNA: competing endogenous RNA
lncRNA: long non-coding RNA
miRNA: micro RNA
mRNA: messenger RNA
ML: Machine Learning
RF: Random Forest
XGB: XGBoost
CSF: Cerebrospinal Fluid
GO: Gene Ontology
BP: Biological Processes
DEG: Differentially Expressed Genes
DBG: Differentially Bound Genes
PGR: Progesterone Receptor
TAD: Topologically Associating Domains
EMT: Epithelial to Mesenchymal Transition

## Supplemental Information Index

- Table S1. Transcript IDs of labour-associated ncRNA identified from literature mining.
- Table S2. Tripartite Motifs comprising of miRNA-lncRNA-TF, matched to database information.
- Table S3. List of DEGs from term and preterm samples analysed with a meta-analysis approach (merged with microarray and RNAseq).
- Table S4. List of DBGs from ChIPseq data between term and preterm samples.
- Table S5. List of TADs with overlapping bipartite RNA combinations.
- Figure S6. Histogram of TAD sizes identified from public myometrial HiC dataset.
- Figure S7. Screenshots of the web server. A) lncRNA-miRNA predictions queried across the 5 models used; B) Graph visualisation of tripartite elements between TF-lncRNA-miRNA.

---

**Fig. 5.**
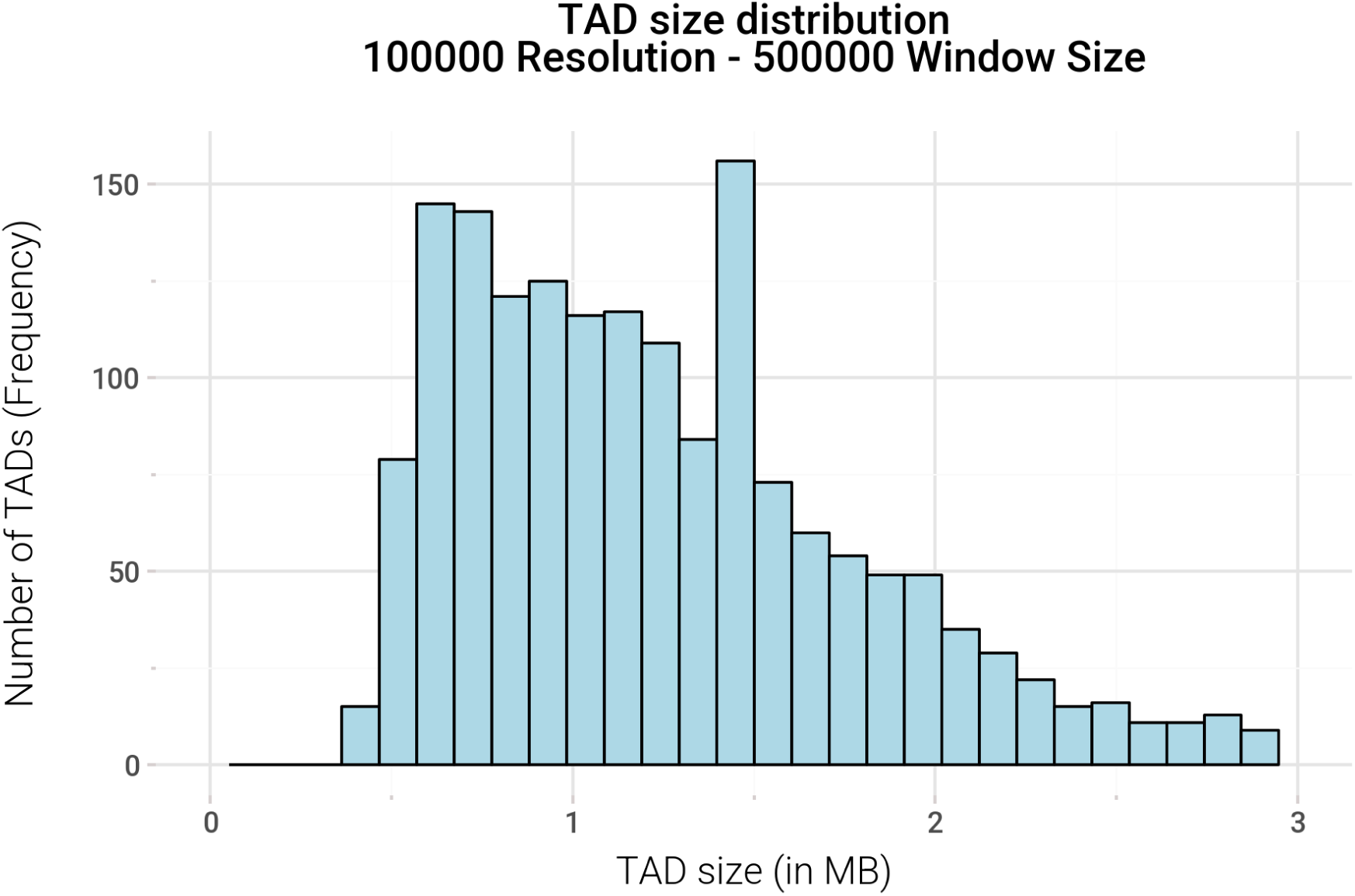
(Supplementary) Histogram of TAD sizes identified from public myometrial HiC dataset.

**Fig. 6.**
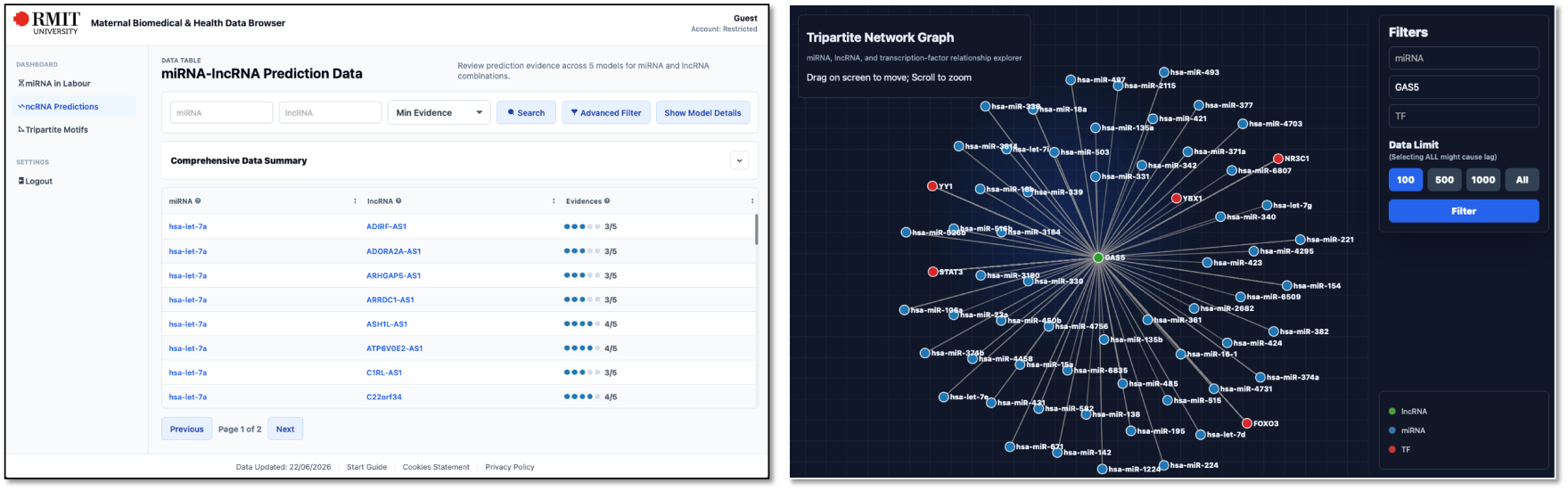
(Supplementary) Screenshots of the webserver. A) lncRNA-miRNA predictions queried across the 5 models used; B) Graph visualisation of tripartite elements between TF-lncRNA-miRNA.

## Methodology

In this work, we used experimentally validated interactions between miRNA and lncRNA as positive data for machine learning (ML) methods to predict potential interactions on new sequences. Four published ML tools and our newly developed ensemble approach were used to predict miRNA-lncRNA interactions (Figures 1A-E; Table 1). The tools were shortlisted by their availability, code access and distinct modelling architectures.

**Fig. 7.**
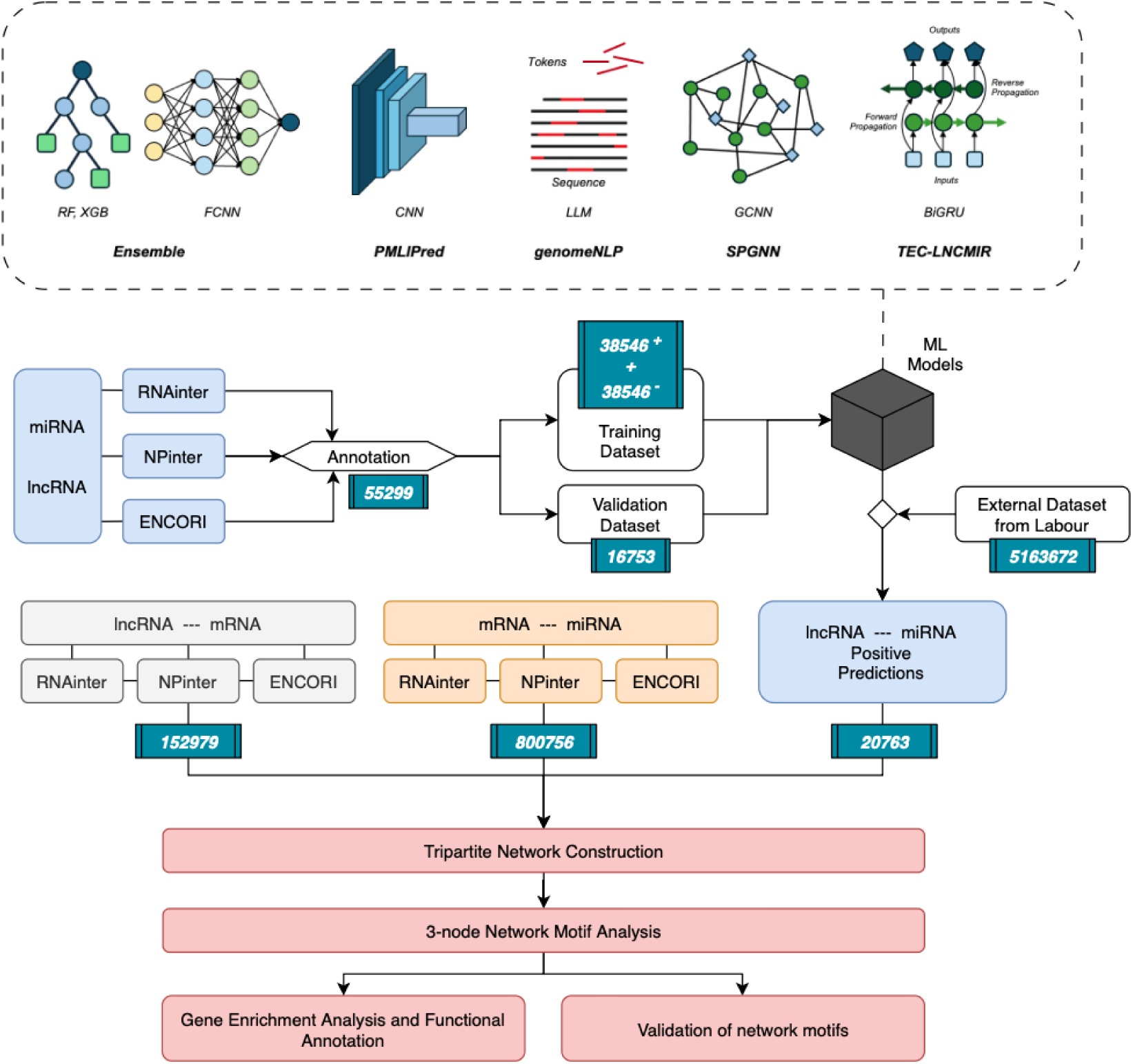
Methodology workflow. Experimentally validated lncRNA-miRNA interactions were used for modelling across 5 architectures, and a separate curated dataset was used for predictions. A tripartite network was constructed with mRNA interactions, followed by network analysis, validation and gene enrichment.

A network of predicted RNA elements and their associated genes is constructed to identify tripartite relationships. The TF-ncRNA links within the network are annotated to gain insight into the regulatory relationships between the elements. The identified tripartite nodes were validated with public omics datasets from myometrial samples.

### Data Collection and Preparation

miRNA (n=2656) and lncRNA (n=191106) sequences were downloaded from miRBase (Release 22.1) and Gencode (Release 42, GRCh38.p14), respectively. lncRNAs for analysis were filtered by the nomenclature overlap of three databases (GENCODE, NONCODE and LNCipedia) [84] and limited to sequence lengths between 200 bp - 11 kbp due to computational constraints. RNA interaction data were collected from STARBase [85], RNAinter [86], and NPinter [87] and curated by species (“*Homo sapiens*”) and supporting experiments (AGO-CLIP). After filtering IDs by available sequences, a total of 55299 miRNA-lncRNA interactions, 800756 miRNA-mRNA interactions and 152979 lncRNA-mRNA interactions were compiled for analysis.

### Data Subset Preprocessing

The model benchmarking was done on the training and the validation datasets. An extra ‘test’ dataset was created by preparing combinations of the labour-associated ncRNA (lncRNA=1427, miRNA=92) with the sequence-available ncRNA elements (miRNA=2656, lncRNA=14930). This data was checked to be mutually exclusive from the training and validation datasets to avoid data leaks and to act as unseen data points for model prediction.

**Table 1.** List of models, their architectures and the data transformation methods used for ncRNA interaction prediction.

| Model Name | Architecture | Feature Engineering | Citation |
| --- | --- | --- | --- |
| Ensemble | Random Forest (RF),<br>Neural Networks (NN),<br>XGBoost (XGB) | K-mer frequency,<br>Doc2Vec embeddings | - |
| genomeNLP | Natural Language processing<br>(NLP) | Tokenisation | [80] |
| SPGNN | Scale-Aware and<br>Global Contextual Network<br>(SGCNet) | Doc2Vec embeddings | [82] |
| PMLI-Pred | Bi-gated Recurrent<br>Neural Network (BiGRU),<br>Random Forest (RF) | One-hot encoding,<br>K-mer frequency | [81] |
| TEC-LncMir | Convolutional Neural Network<br>(CNN) | One-hot encoding | [83] |

### Model Selection

Multiple tools have been published to predict RNA:RNA interactions, and they rely on sequence information or expression data with appropriate modelling methods to predict interactions [69]. After a brief literature search, multiple deep learning architectures such as CNNs, RNNs, and graph networks for RNA:RNA interaction prediction were identified, and Python-based tools were investigated for implementation. We implemented 4 published RNA:RNA interaction tools – PMLIPred [81], genomeNLP [80], SPGNN [82] and TEC-LncMir [83]. TEC-LncMir uses RNA sequences to create one-hot feature embeddings and models them using a CNN architecture. genomeNLP is a genomic large language model (LLM) based pipeline that utilises nucleotide sequence data embedded model called genomicBERT for classification tasks. SPGNN is a graph-based model that creates a network using each RNA as a node, and the interaction scores between RNA types are mapped as the edges. The edges are weighted based on the sequence embeddings calculated for the nodes, and potential interactions are calculated by training on the network. PMLIPred uses a fuzzy late-fusion architecture made of 2 models – BiGRU and Random Forest (RF). The sequence k-mer features are modelled using RF, and the BiGRU architecture is trained on RNA sequence embeddings. The output from both these models is merged as a weighted average. Additionally, to check the predictive power of shallow ML methods, an ensemble model comprising 3 different shallow ML models - Random Forest (RF), XGBoost (XGB) and Fully Connected Neural Network (FCNN) was also implemented for modelling and predicting ncRNA interactions.

### Feature Engineering

**Table 2.** Summary of feature generation and dimensions of generated embeddings. Category 1 features were combined together to create a 172-dimensional vector. This was merged with one type of category 2 for ensemble modelling.

| Category | Features | Length | Type | Transformation | Models Used |
| --- | --- | --- | --- | --- | --- |
| 1 | 1-mer | 4 | Text | Counts | RF, XGB, NN |
| 1 | 2-mer | 16 | Text | Counts | RF, XGB, NN |
| 1 | 3-mer | 64 | Text | Counts | RF, XGB, NN |
| 1 | GC% | 1 | Numerical | - | RF, XGB, NN |
| 1 | Length | 1 | Numerical | - | RF, XGB, NN |
| 2A | Doc2Vec (3-mer) | 128 | Text | Embedding | RF, XGB, NN |
| 2B | Doc2Vec (6-mer) | 128 | Text | Embedding | RF, XGB, NN |
| 2C | Doc2Vec (3-mer) | 256 | Text | Embedding | RF, XGB, NN |
| 2D | Doc2Vec (6-mer) | 256 | Text | Embedding | RF, XGB, NN |

The complete corpus of lncRNA and miRNA sequences was independently converted into feature embedding models using the Doc2Vec [88] method. The RNA:RNA interaction datasets were mapped to their sequences and run through 2 feature generation methods: K-merisation and Doc2Vec embedding. Table 2 summarises the different features generated for ensemble modelling.

### ML Modelling

The training data (n=77092) was run through all 5 models after appropriate feature processing. The ensemble model was trained with multiple combinations of the features (categories in Table 2) and the ML models, and a hard-voting classifier was used to compile the predictions. The best ensemble combination was selected based on the calculated accuracy and AUROC scores. For genomicBERT, a token file of vocabulary size 4000 was used to generate the embeddings for the training data. Each of the 5 models was trained independently and was used to predict on the test dataset after training. The prediction results were compiled from all models, and positive predictions from 4 or more models were used for network analysis.

### Network Motif Analysis

The predicted list of lncRNA-miRNA bipartite interactions was merged with the compiled mRNA-lncRNA and mRNA-miRNA interactions. The Python package networkX (version 3.3) was used to identify 3-node cliques (which we refer to as tripartite motifs) from the compiled bipartite information. The resultant tripartite motifs were converted to a simple interaction format (SIF) file for network visualisation and analysis in Cytoscape (version 3.10.3).

### Gene Enrichment Analysis

A custom Python script was used to classify the RNAs within the tripartite motifs into their respective categories (lncRNA, miRNA and mRNA) for gene set enrichment analysis (GSEA). GProfiler [27], STRING [89], and miEAA [90] are online tools used for functional annotation across categories (mRNA, lncRNA, miRNA). The enrichment results for each category of RNA were sorted by their gene ratio, and the top 15 adjusted p-values were plotted against the term names. Additionally, the tripartite motifs containing TFs were isolated and analysed using STRING. MCL clustering algorithm [91] with an inflation parameter of 3 was used to identify functionally linked nodes and to annotate terms across various categories such as biological processes, molecular function, cellular compartmentalisation, tissues and molecular pathways.

**Fig. 8.**
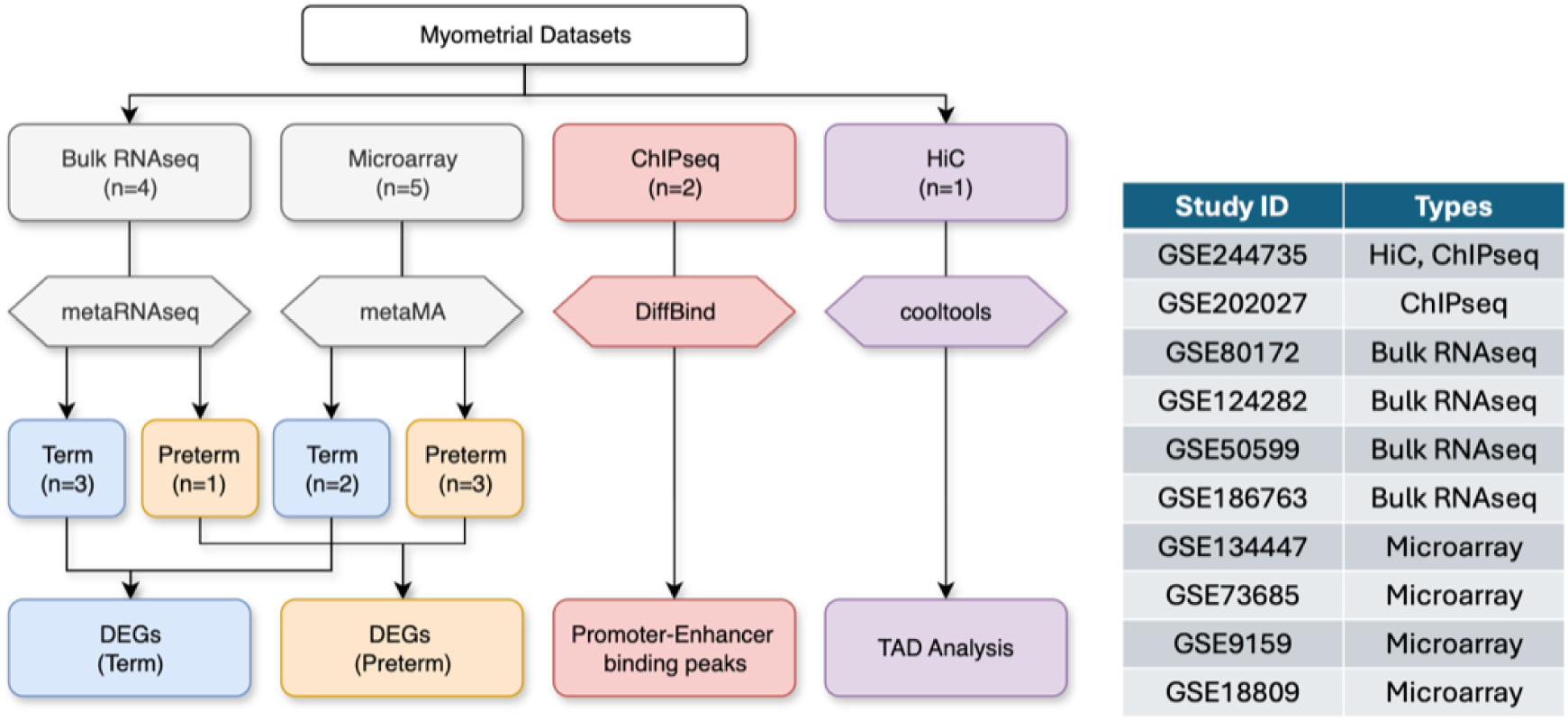
List of public, myometrial datasets and workflows used for validation of motifs.

### Genomic Localisation Analysis

The network nodes were mapped to their genomic loci using the GTF file downloaded from GENCODE (GRCH38.p14). The observed frequencies of the network nodes for TF, lncRNA and miRNA were calculated independently by type against the total frequency of the gene type across all chromosomes. The expected frequencies of TF, lncRNA and miRNA were computed using the total genes present in each chromosomes against the genome. A ratio of observed/expected frequency was plotted to identify the degree of involvement across each chromosome.

### Network Validation

Public databases were searched to identify studies on pregnant myometrial samples (Figure 8). The inclusion criteria were limited to myometrial tissues, with flexibility on term or preterm status. Samples with co-morbidities or other disease states (e.g. chorioamnionitis, pre-eclampsia) were excluded from the analysis. A meta-analysis approach using the R packages ‘metaMA’ [92] for microarray data and ‘metaRNAseq’ [93] for Bulk RNAseq data was implemented to analyze and identify DEGs (Figure 8). The merged results were mapped to the network tripartite motifs to infer their expression patterns in labouring and non-labouring statuses. ChIP-seq and HiC datasets from myometrial tissues were used to visualize the binding peaks and chromatin conformation, respectively, for the DEG tripartite motifs.

#### Transcriptomics analysis and meta-analysis

Microarray data was downloaded from NCBI and Cell Intensity File (CEL) files were converted to matrix files using the Affy and limma packages. The matrix files were merged and analyzed using the metaMA package. Bulk RNAseq FASTQ files were downloaded and processed with the Nextflow nf-core/rnaseq pipeline (version 3.14.0). The expression matrix files were analyzed together with the metaRNAseq package. Genes with *abs*(*logFC*) *>*= 1.5 and *p* − *value <* 0.05 were selected as differentially expressed genes, and compared to the network nodes.

#### ChIP-seq analysis

Myometrial ChIP-seq FASTQ data were downloaded from NCBI and analyzed using the Nextflow nf-core/chipseq pipeline (version 2.1.0). The peak calling and differential binding analysis was done using MACS2 and Homer. The differentially bound genes (DBGs) were compiled for each available protein (histones and TF) and cross-referenced with the motif network. The binding peaks of the DBGs were visualized using pyGenomeTracks (version 3.1.2)[94].

#### HiC analysis

Myometrial HiC data (GSE244735) was downloaded from NCBI and analyzed using the runHiC pipeline (version 0.8.7). The chromosome contact matrix was processed at 100Kb resolution, and the contact map was visualised using the cooltools python package (version 0.7.1) [95]. The “insulation()” function in cooltools was used to identify the TAD boundary sites using a sliding window size of 500 Kb. The motifs were mapped to TAD regions based on their genomic loci. Intra-chromosome distances between motif loci was calculated using the matrix probability scores and the formula suggested by Shi *et al.* [96]. The expected probability as a function of genomic separation was calculated using the in-built “expected_cis()” function in cooltools. The calculated and expected distance scores were analysed using a 2-sided Kolmogorov-Smirnov (KS) statistical testing [2].

### Database Development

A public database has been constructed and made available to browse the results compiled in this study. The lncRNA-miRNA predicted interactions and the tripartite motifs with TFs can be viewed in a tabular format with custom query options. An interactive graph network can be visualised for the tripartite motifs (Supplementary Figure 6).

